# Reprogrammed peptidoglycan elongation reveals plasticity in bacterial growth modes

**DOI:** 10.64898/2026.04.28.721452

**Authors:** Carolina Basurto De Santiago, Isabella S. Lin, Beiyan Nan

## Abstract

Most bacteria are enclosed by a peptidoglycan (PG) cell wall that must be expanded for growth. In rod-shaped species, PG elongation is spatially organized in a species-specific manner, occurring either at the cell poles or along the lateral wall. MreB filaments organize the Rod machinery and are typically required for dispersed, nonpolar PG elongation but are dispensable for polar growth. Whether elongation modes are inherently fixed or can be reprogrammed remains unclear. *Escherichia coli* and *Myxococcus xanthus* both elongate PG in a dispersed, nonpolar fashion. Here, we show that heterologous expression of *M. xanthus* MreB in *E. coli* relocalizes native MreB to the cell poles, thereby redirecting the Rod machinery and PG elongation to polar sites. Moreover, direct targeting of the Rod synthase PBP2 to the poles is sufficient to drive polar PG elongation in *E. coli* while preserving rod shape. This reprogrammed growth mode bypasses the requirement for MreB filaments, highlighting a plasticity of the Rod system that suggests polar elongation may have emerged through the evolutionary loss of MreB.

---

Most bacterial cells utilize the peptidoglycan (PG) cell wall as a major stress-bearing structure that determines cell shape and prevents cells from lethal rupture caused by high osmotic pressure [1, 2]. To grow in size, bacteria expand their existing PG structures using distinct patterns. In rod-shaped bacteria, two main modes of PG elongation are observed: polar elongation, where new cell wall material is added at one or both poles, and dispersed nonpolar elongation, which involves a more uniform insertion of PG along the lateral sides of the cell. These elongation strategies are typically species-specific, with each organism adopting either one mode or a combination of both [3, 4]. Although most well-studied rod-shaped bacteria exhibit dispersed elongation, polar elongation is notably characteristic of Gram-negative *Rhizobiales* and Gram-positive *Actinomycetales* [5-7].

PG of both Gram-positive and negative bacteria consists of glycan strands that are covalently crosslinked by short peptides [8, 9]. PG precursors are synthesized in the cytoplasm and flipped across the cytoplasmic membrane, where PG synthases, including glycosyltransferases (**GTases**) that polymerize them into glycan strands, which are then crosslinked into PG network by transpeptidases (**TPases**). In most rod-shaped bacteria, two synthase systems assemble the PG during cell elongation, the Rod system and class A penicillin-binding proteins (**aPBPs**) [2, 10, 11]. The Rod system is a multiprotein complex that includes RodA, a shape, elongation, division, and sporulation (**SEDS**) family GTase; PBP2, a TPase of the class B penicillin-binding protein (**bPBP**) family; and the actin homolog MreB. This system defines rod shape and drives the majority of PG expansion during vegetative growth [11-13]. In contrast, aPBPs that possess both GTase and TPase activities do not depend on MreB, and the absence of single aPBPs rarely abolishes cell survival or rod-like morphology [12, 14-17].

MreB is essential for rod shape in the cells that elongate in dispersed nonpolar manner [13]. MreB polymerizes into antiparallel double filaments on the inner leaflet of the cytoplasmic membrane [18, 19]. Sensitive to membrane curvature, MreB filaments guide RodA and PBP2 to the cylindrical regions of the cell envelope, where PG elongation occurs [13, 20-23]. Depleting MreB or inhibiting MreB polymerization using A22, a specific inhibitor, causes cells to lose rod shape in a wide range of model organisms, including *Escherichia coli* [24, 25], *Caulobacter crescentus* [26], *Pseudomonas aeruginosa* [27], *Bacillus subtilis* [28], *Myxococcus xanthus* [29], etc. Conversely, MreB is either absent or nonessential in the bacteria that elongate at the poles [30]. Instead, these organisms repurpose cell division related proteins, such as FtsA and FtsZ in *Rhizobiales* and DivIVA in *Actinomycetales* to localize PG synthases to cell poles [31, 32]. Recent studies revealed that closely related bacterial species in the family *Caulobacteraceae* can adopt different PG elongation modes [3] and that Streptomyces can use both polar and dispersed elongation modes in a special growth stage called exploratory growth [4]. Nonetheless, it remains uncertain whether the PG elongation mode in rod-shaped bacteria can be entirely reprogrammed without compromising cell morphology.

*E. coli* (γ-proteobacterium) and *M. xanthus* (long assigned to δ-proteobacteria but recently reclassified into the newly established phylum Myxococccota [33]) both elongate their PG in dispersed manner using their respective Rod systems [17, 20, 34-36]. In this study, we found that the heterogeneous expression of *M. xanthus* MreB (**MreB**_**Mx**_) in wild-type *E. coli* caused the native MreB (**MreB**_**Ec**_) to form aggregates at cell poles and thereby redirect the Rod synthases to elongate PG at poles. Building on this observation, we further demonstrated that directly targeting PBP2 to cell poles is sufficient to switch *E. coli* PG elongation to poles. In both cases, the reprogrammed polar PG elongation maintains rod shape independent of MreB filaments. Given that MreB is an ancient protein believed to have existed prior to the emergence of *Rhizobiales* and *Actinomycetales* [37], it is plausible that polar growth may have evolved from dispersed elongation as a consequence of MreB loss.

## Results

### Heterologous expression of MreB_Mx_ in *E. coli* causes MreB_Ec_ to aggregate at cell poles

While cloning the *M. xanthus mreB* (***mreB***_**Mx**_) gene into the plasmid pMAT3 [38] using the *E. coli* DH5α strain, we found that *E. coli* cells harboring the pMAT3-*mreB*_Mx_ plasmid grew slower and displayed dark aggregates near their cell poles under bright field microscopy (**Fig. 1a**). To determine whether this phenomenon is specific to the DH5α strain, we transformed the pMAT3-*mreB*_Mx_ plasmid into wild-type *E. coli* strains MG1655 and BW25113 and observed similar aggregates at cell poles (**Fig. 1a**). In contrast, the empty pMAT3 vector did not cause polar aggregation (**Fig. 1a**, insets). To maintain consistency, we performed all subsequent experiments in MG1655-derived strains.

**Fig. 1.**
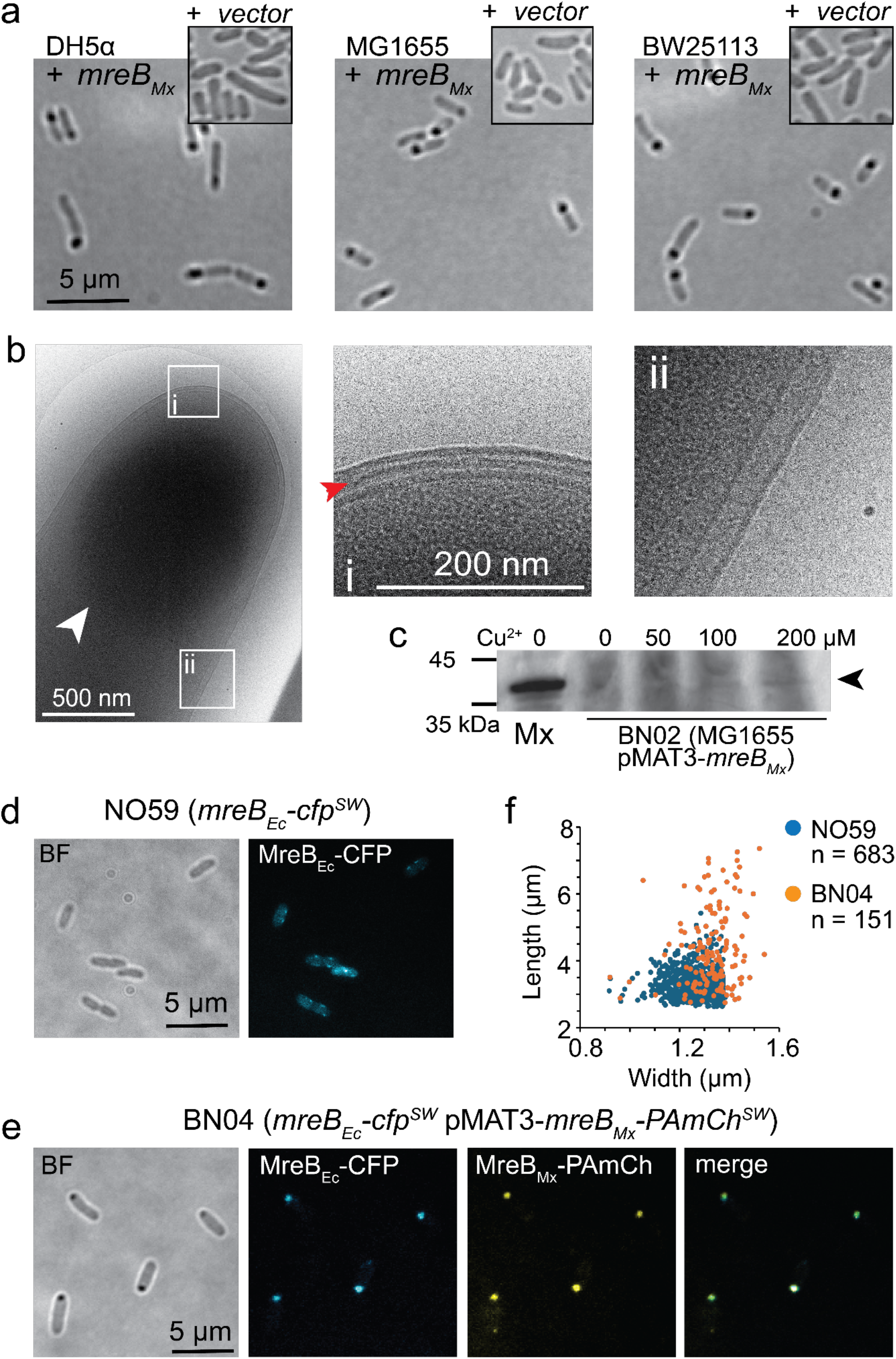
Heterologous expression of MreB_Mx_ in *E. coli* causes MreB_Ec_ to aggregate at cell poles. **a)** Heterologous expression of MreB_Mx_ in *E. coli* strains DH5α, MG1655, and BW25113 causes *E. coli* cells to form dark aggregates at cell poles. Insets show the morphology of cells carrying the empty pMAT3 vector. **b)** CryoEM images of an *E. coli* MG1655 cell that expresses MreB_Mx_. White arrow points to the polar aggregate. In the cells with polar aggregates, PG thickens at cell poles (inset i, red arrow), whereas PG is rarely visible in nonpolar regions (insert ii). **c)** MreB_Mx_ expresses at very low levels, even when induced by high concentration of CuSO_4_. The native expression of MreB_Mx_ from the same number of *M. xanthus* (Mx) cells is shown in the first lane as a reference. Black arrow points to the bands detected by an anti-MreB_Mx_ antibody. **d)** MreB_Ec_-msfCFP^SW^ forms dispersed foci in the NO59 strain. **e)** In BN04 (NO59 pMAT3-*mreB*_*Mx*_*-PAmCherry*^*SW*^) cells, instead of forming filaments, MreB_Mx_ and MreB_Ec_ colocalize in polar aggregates. **f)** While BN04 strain retains rod shape, expressing MreB_Mx_ increases both the length and width of cells. BF, bright field.

Under a cryogenic electron microscope (**cryoEM**), these aggregates appeared as dense spheres that lack clear boundaries (**Fig. 1b**), similar to the inclusion bodies frequently observed in *E. coli* cells that express recombinant proteins [39]. Remarkably, in cells with polar aggregates, we observed dark densities from the PG layers at the cell poles (**Fig. 1b i**), whereas PG was rarely visible in nonpolar regions (**Fig. 1b ii**). The correlation between polar aggregates and thickened PG suggests that these aggregates may enhance local PG synthesis.

The pMAT3 plasmid, which carries the ColE1 origin, was designed for gene expression under the control of a copper-inducible promoter [38]. However, cells carrying the pMAT3-*mreB*_*Mx*_ plasmid formed aggregates even in the absence of added copper. We reasoned that the leaky expression from the promoter was unlikely to produce MreB_Mx_ at levels high enough to form inclusion bodies. To assess how MreB_Mx_ expression influences aggregate formation in *E. coli*, we performed immunoblotting. As shown in **Fig. 1c**, the highly sensitive anti-MreB_Mx_ antibody [40] did not detect significant MreB_Mx_ expression in the absence of copper, indicating that the leaky expression was extremely low and that the antibody did not cross-react with MreB_Ec_ (**Fig. 1c**). Even at CuSO_4_ concentrations as high as 200 µM, MreB_Mx_ expression was only modestly induced in *E. coli*, to less than 5% of its native expression level in *M. xanthus* (**Fig. 1c**). Collectively, these findings suggest that even minimal levels of MreB_Mx_ are sufficient to initiate aggregate formation. In this context, MreB_Mx_ acts not as a primary structural component of the aggregates, but rather as a trigger for their formation. Therefore, in all subsequent experiments, we expressed MreB_Mx_ from pMAT3 without the addition of copper. In this condition, among 799 cells we analyzed, 68.1% contained one aggregate, 3.4% contained two, while 28.5% did not show significant aggregates. We reason that this variability may stem from the heterogeneity of the leaky MreB_Mx_ expression among individual cells and the limited sensitivity of bright field microscopy.

Given that MreB_Mx_ and MreB_Ec_ share 63.3% amino acid sequence identity [41], we hypothesized that even low levels of MreB_Mx_ could interfere with MreB_Ec_ filament assembly, causing MreB_Ec_ to form inclusion body-like aggregates. In the MG1655-derived *E. coli* strain NO59 that expresses an internal monomer super-folder cyan fluorescence protein (**msfCFP**^**SW**^)-labeled MreB_Ec_ from its native locus and promoter [42], short MreB_Ec_ filaments appear as foci on cell peripherals (**Fig. 1d**). To test our hypothesis, we generated the strain BN04 by expressing an internal photoactivatable mCherry (**PAmCherry**^**SW**^)-labeled MreB_Mx_ that is functional in *M. xanthus* [29] in NO59 using the pMAT3 vector. In bright field images, BN04 cells showed polar aggregates (**Fig. 1e**) similar to those observed in the MG1655 cells expressing the unlabeled MreB_Mx_ (**Fig. 1a**). Thus, introducing fluorescent tags into both MreB homologs did not affect aggregate formation. When exposed to 405-nm excitation (0.2 kW/cm^2^) for 2 s, the majority of MreB_Mx_-PAmCherry^SW^ was photoactivated [29]. In the cells that contained aggregates, the emission signals of both MreB_Mx_ and MreB_Ec_ colocalized with the polar aggregates (**Fig. 1e**). Notably, MreB_Ec_ filaments were never observed (**Fig. 1e**), indicating that heterologous expression of MreB_Mx_ in *E. coli* causes the majority of MreB_Ec_ to aggregate at cell poles.

### Polar aggregation of MreB_Ec_ causes *E. coli* to elongate PG at cell poles

Polar aggregation of MreB_Ec_ clearly affected PG elongation: expression of MreB_Mx_-PAmCherry^SW^ increased the generation time from 41.2 ± 3.0 min (n = 4) of NO59 to 127.6 ± 8.6 min (n = 3) of BN04 (**Fig. S1**) and led to a 33.11% and 5.70% increase in cell length and width, respectively (**Fig. 1f**). However, cells still retained rod shape.

To visualize newly synthesized PG, we used BODIPY-FL 3-amino-D-alanine (**BADA**), a fluorescently labeled D-alanine analog that incorporates into the PG scaffold during elongation [43, 44]. To visualize PG assembly on the entire cell surface, we used fluorescence microscopy to visualize a ∼200-nm thick longitudinal section close to the coverslip under highly inclined and laminated optical sheet (HILO) illumination [29, 35, 45, 46] (**Fig. 2a**). In the NO59 strain, the incorporation of BADA appeared in a dispersed pattern, aligning with the distribution of MreB_Ec_ filaments. Notably, the absence of BADA at the cell poles reinforces the nonpolar elongation mode (**Fig. 2a, b**). In contrast, in 71.81% (n = 674) of the BN04 cells, PG elongation occurred exclusively at cell poles, colocalizing with the aggregates that contained both MreB_Ec_ and MreB_Mx_ (**Fig. 2d, e**).

**Fig. 2.**
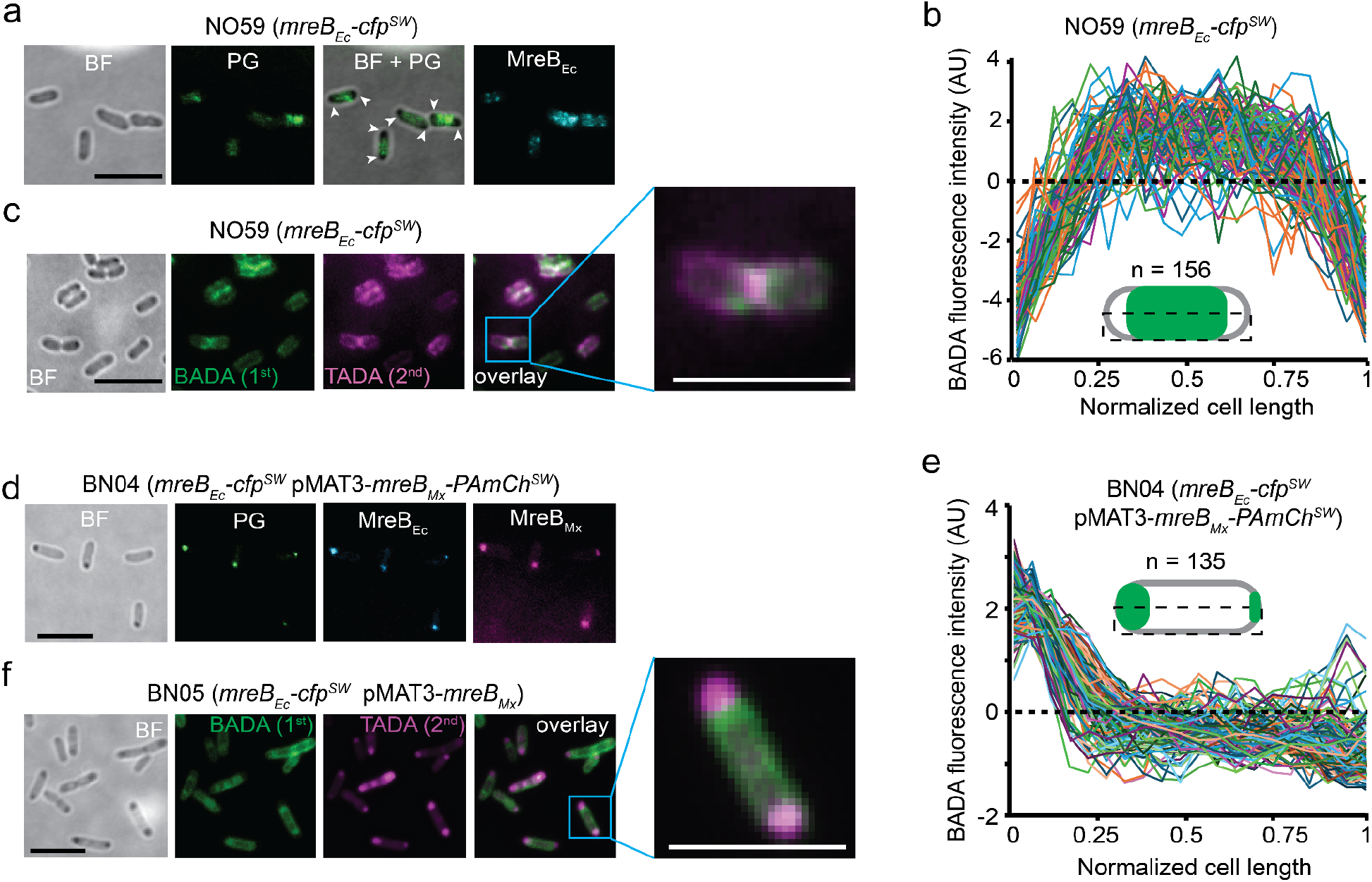
Heterologous expression of MreB_Mx_ causes *E. coli* to elongate PG at cell poles. **a)** NO59 cells elongate PG at nonpolar regions in a dispersed manner, consistent with MreB_Ec_ localization pattern. PG elongation was labeled by BADA. White arrows point to cell poles. **b)** Quantitative analysis of PG elongation across 156 NO59 cells. Cell lengths were normalized to 1 and BADA fluorescence intensity profiles were standardized using Z-score normalization (0 indicates the mean value for each cell). Insets here and in panel **e** illustrate patterns of PG elongation, with newly synthesized PG shown in green. Imaged cell regions are highlighted by dotted-line boxes. **c)** PG elongation mode in NO59 cells visualized by pause-chase sequential labeling with BADA and TADA, where both dyes were incorporated in a dispersed, nonpolar pattern. **d)** BN04 cells that express MreB_Mx_-PAmCherry^SW^ elongate PG at poles, consistent with the localization patterns of both MreB_Ec_ and MreB_Mx_. PG elongation was visualized by BADA. **e)** Quantitative analysis of PG elongation across 135 BN04 cells. For each cell, its BADA fluorescence intensity profile was measured starting from the only or brighter cell pole. **f)** Elongation mode in BN05 cells visualized by pause-chase sequential labeling with BADA and TADA. While the new growth (TADA) occurred at cell poles, the old growth (BADA) receded to nonpolar regions. BF, bright field. Scale bars, 5 μm.

To further validate this observation, we conducted a pulse-chase experiment to visualize PG elongation using sequential incorporation of two fluorescent D-alanine analogs. For the NO59 strain, we first stained PG elongation using BADA for 15 min, washed BADA out, then switched to the second dye, TAMRA 3-amino-D-alanine (**TADA**) for another 15 min. Consistent with its nonpolar elongation mode, both dyes were incorporated in the same dispersed pattern into lateral cell surfaces (**Fig. 2c**). To facilitate pause-chase labeling, we constructed a BN05 strain that expressed unlabeled MreB_Mx_ in the NO59 background and visualized PG elongation using BADA and TADA sequentially. While the second dye, TADA, was incorporated exclusively at the cell poles, the first dye, BADA, receded to subpolar regions (**Fig. 2f**), indicating that newer PG elongation occurs exclusively at poles. These results suggest that some MreB_Ec_ molecules in polar aggregates might be recruiting the Rod enzymes to poles and thus redirecting *E. coli* growth to a polar elongation mode.

### The Rod system carries out the reprogrammed, polar PG elongation

To determine whether aPBPs or the Rod system drives polar elongation when MreB_Ec_ aggregates at cell poles, we inhibited their activities by moenomycin (10 µg/ml) and mecillinam (100 µg/ml), respectively. While moenomycin blocks all aPBPs [47], mecillinam specifically inhibits PBP2 in the Rod system [48]. Under both treatments, the NO59 cells lost their rod shape but continued to incorporate BADA without a recognizable pattern (**Fig. 3a**), suggesting that although both aPBPs and the Rod system are required for rod shape in *E. coli*, either system is sufficient for PG elongation, albeit in disorganized manners. Notably, MreB_Ec_ appeared diffusive under both conditions (**Fig. 3a**), indicating extensive disassembly of MreB_Ec_ filaments, for which the mechanisms remain to be investigated.

**Fig. 3.**
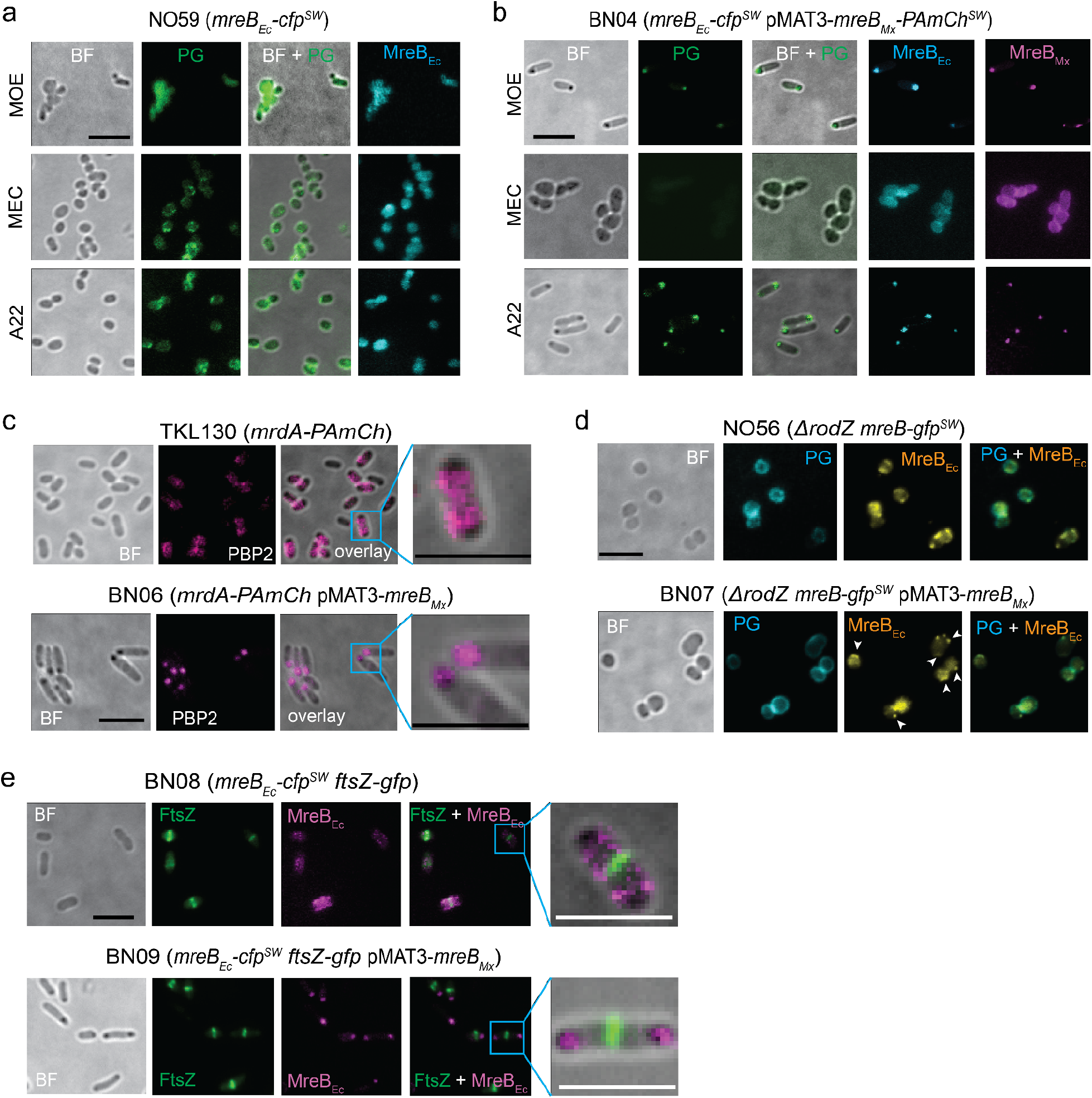
The relocalized Rod system carries out the reprogrammed, polar PG elongation. **a)** In the absence of heterologous MreB, *E. coli* cells elongate PG using both the Rod system and aPBPs. Neither moenomycin (MOE, 10 μg/ml) that inhibits all aPBPs, mecillinam (MEC, 100 μg/ml) that inhibits PBP2 in the Rod system, or A22 (10 μg/ml) that inhibits the polymerization of MreB filaments and thus disrupts the assembly of Rod complexes, abolishes PG elongation completely. PG elongation was visualized using BADA. **b)** *E. coli* cells expressing MreB_Mx_ solely rely on the Rod system for polar PG elongation, as MEC alone is sufficient to abolish PG elongation. Such polar PG elongation is resistant to A22, suggesting that polar localized Rod enzymes elongate PG independent of MreB filaments. PG elongation was visualized using BADA. **c)** Heterologous expression of MreB_Mx_ recruits *E. coli* PBP2 to cell poles. **d)** In the absence of RodZ, MreB_Ec_ still forms aggregates at pole-like locations (white arrows) but fails to induce focused PG growth. PG elongation was visualized using BADA. **e)** The reprogrammed, polar PG elongation does not depend on the divisome as FtsZ does not colocalize with polar MreB aggregates.

Under moenomycin treatment, the BN04 cells that expressed MreB_Mx_ retained their rod shape and continued PG elongation at their poles, with both MreB_Ec_ and MreB_Mx_ remained at the poles (**Fig. 3b**). Thus, focused polar growth fully bypassed the requirement for aPBPs. In contrast, mecillinam inhibited PG growth and abolished rod shape in BN04 cells (**Fig. 3b**). As mecillinam is sufficient to block polar PG elongation, the Rod system is the sole player that drives polar PG elongation. Interestingly, the expression of MreB_Mx_ did not cause MreB_Ec_ to aggregate in the presence of mecillinam (**Fig. 3b**). As mecillinam blocks MreB_Ec_ filament assembly (**Fig. 3a**), our observation suggests that MreB_Mx_ may only cause assembled MreB_Ec_ filaments to aggregate.

To drive focused PG elongation at cell poles, the Rod system enzymes RodA (GTase) and PBP2 (TPase) must localize to the poles. The TKL130 strain, derived from MG1655, expresses a PAmCherry-labeled PBP2 from its native locus and promoter [49]. Under a 405-nm excitation (0.2 kW/cm^2^, 2 s) that activates the majority of PAmCherry, PBP2 localized in a dispersed pattern (**Fig. 3c**). To test if PBP2 switches its localization to cell poles while the Rod system drives polar PG elongation, we generated the BN06 strain by introducing the pMAT3-*mreB*_Mx_ plasmid into the TKL130 background. Similar to our observation on BN04 and BN05 cells, heterologous expression of MreB_Mx_ caused BN06 cells to form dark polar aggregates under bright-field microscopy, which colocalized with bright PBP2 foci (**Fig. 3c**).

To further test if MreB_Ec_ aggregates recruit the Rod polymerases to cell poles, we investigated whether they still trigger polar PG elongation in the absence of RodZ, the connector between MreB_Ec_ and Rod enzymes [42]. The NO56 cells that expressed MreB_Ec_-msfGFP^SW^ in a *ΔrodZ* background were near spherical (**Fig. 3d**). Consistent with the disconnection between MreB_Ec_ and Rod enzymes, while MreB_Ec_ still formed filaments, HCC-amino-D-alanine (**HADA**), another fluorescent D-amino acid, was incorporated near homogeneously on the entire cell surface (**Fig. 3d**). We introduced pMAT3-*mreB*_*Mx*_ into the NO56 background to generate the BN07 strain. Similar to the parental NO56 cells, BN07 cells were near spherical, with imperfect local curvatures that mimicked cell poles (**Fig. 3d**). Strikingly, while MreB_Mx_ expression caused MreB_Ec_ to aggregate at such pole-like locations, these aggregates neither induced focused HADA incorporation, nor restored rod shape (**Fig. 3d**). Therefore, in the absence of RodZ, MreB_Ec_ aggregates fail to induce focused PG elongation. Taken together, in the reprogrammed, polar-elongating cells (BN04, BN05, and BN06), MreB_Ec_ aggregates recruit the Rod polymerases to poles through RodZ.

Besides the Rod system and aPBPs, a third machinery, the divisome, specifically assembles PG at the division site. To further confirm that the divisome does not significantly contribute to polar PG elongation, we visualized the localization of FtsZ, the cytoskeletal element that orchestrates the PG synthases in the divisome, by expressing a GFP-labeled FtsZ as merodiploid [50] in the NO59 (*mreB*_*Ec*_*-cfp*^*sw*^) and BN05 (*mreB*_*Ec*_*-cfp*^*sw*^ pMAT3-*mreB*_*Mx*_) backgrounds. In both strains, FtsZ-GFP forms bright bands in the center of cells, marking the current or future division sites (**Fig. 3e**). Notably, in the BN05 background, FtsZ did not colocalize with the polar MreB_Ec_ aggregates. Thus, the divisome does not contribute to the polar PG elongation. On the other hand, the reprogrammed PG elongation does not change the mode of cell division. Taken together, the polar localized Rod system is both sufficient and necessary for focused PG elongation at poles.

### Targeting PBP2 to cell poles is sufficient to drive polar PG elongation

To determine whether *E. coli*’s elongation mode could be reprogrammed by directly targeting Rod polymerases to cell poles, we modified the PopZ-Linked Apical Recruitment (POLAR) system [51] to express a PopZ-H3H4-PBP2 fusion *in trans* in the NO59 strain with an arabinose-inducible promoter using the pHCL149 plasmid [51]. In this fusion, PopZ from *C. crescentus* has been proven to localize to *E. coli* cell poles while the H3H4 homo-oligomerization domain promotes targeting PBP2 to polar PopZ clusters [52]. Prior to arabinose induction, 95.93% (n = 123) of cells exhibited dispersed PG elongation and MreB_Ec_ distribution (**Fig. 4a**). Upon induction, 92.06% (n = 126) of cells displayed polar PG elongation that colocalized with MreB_Ec_ aggregates (**Fig. 4a**). These results indicate that targeting a critical amount of PBP2 to cell poles is sufficient to shift *E. coli* growth to polar elongation. In addition, the fact that MreB_Ec_ forms polar aggregates following PBP2 indicates that PBP2 can regulate the localization of MreB.

**Fig. 4.**
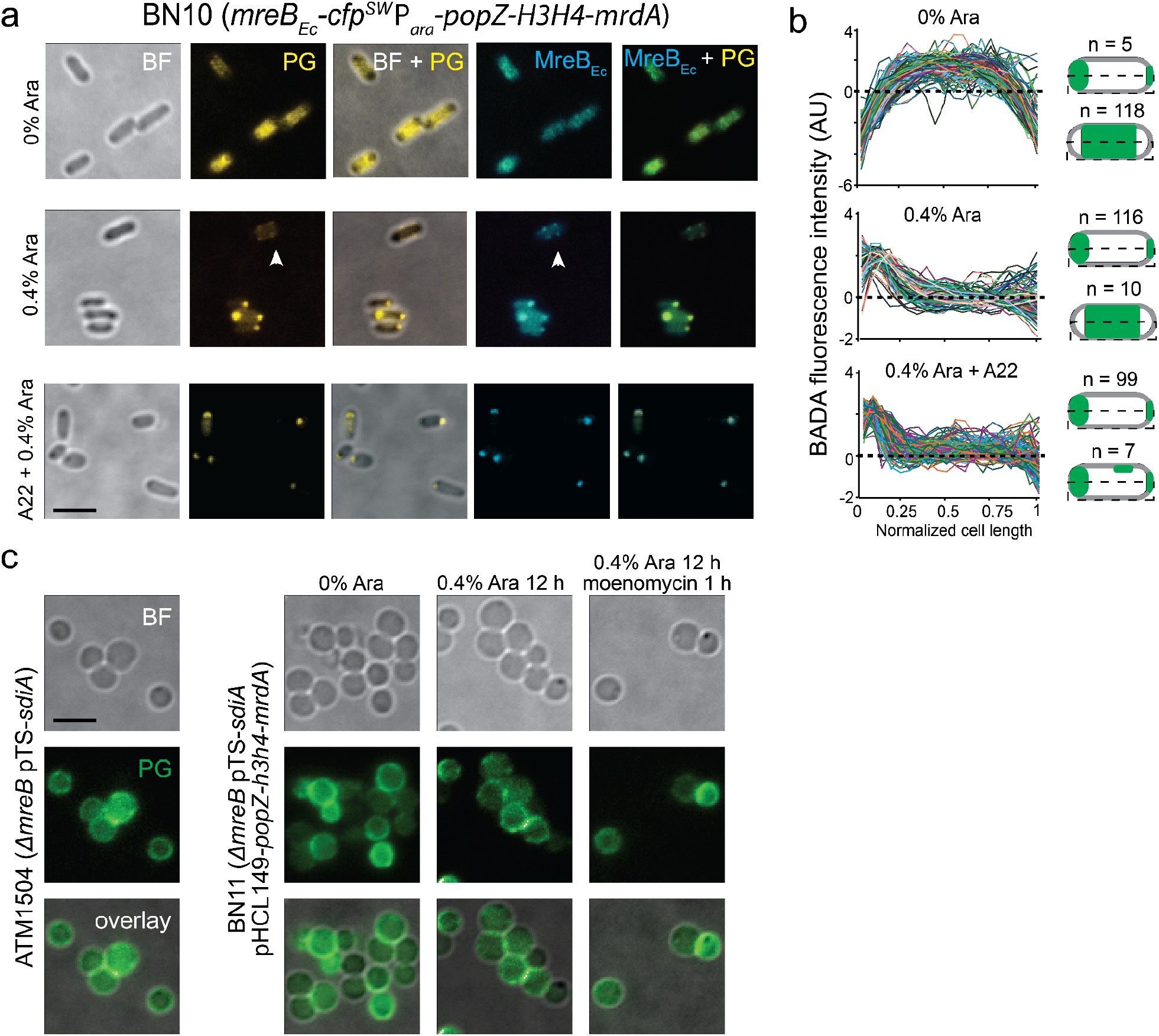
Targeting PBP2 to cell poles is sufficient to drive polar PG elongation. **a)** When PBP2 (encoded by *mrdA*) is ectopically expressed with a polar-targeting tag (PopZ-H3H4) by an arabinose-inducible promoter under 0.4% arabinose (Ara), MreB_Ec_ aggregates at cell poles and cells switch to polar PG elongation and this reprogrammed growth mode is resistant to A22 (10 μg/ml). The white arrow points to a cell that still elongated PG at nonpolar sites. **b)** Quantitative analysis of PG elongation. Cell lengths were normalized to 1 and BADA fluorescence intensity profiles were standardized using Z-score normalization (0 indicates the mean value for each cell). Insets illustrate patterns of PG elongation, with newly synthesized PG shown in green. The numbers of cells that adopt each elongation pattern were presented above the insets. Imaged cell regions are highlighted by dotted-line boxes. **c)** Expressing pole-targeting PBP2 in the absence of MreB_Ec_ neither induces focused PG elongation (labeled by BADA) nor restores rod-shape, despite that Rod enzymes still incorporate BADA. aPBPs were inhibited by moenomycin (MOE, 4 μg/ml) for 1 h before adding BADA for 15 min. Scale bars, 5 μm.

### Polar PG elongation does not require MreB filaments

Because cells continue to elongate PG at their poles despite most MreB_Ec_ being sequestered in aggregates, polar PG elongation requires few, if any, MreB_Ec_ filaments. To clarify the roles of MreB filaments in polar elongation, we first treated cells with A22 that inhibits MreB polymerization [18, 53]. In the NO59 cells that expressed MreB_Ec_-msfCFP^SW^, A22 (10 µg/ml) abolished rod shape and dispersed MreB_Ec_ foci within 1 h, albeit cells continued to incorporate BADA, likely through aPBPs (**Fig. 3a**). These results confirm numerous reports that MreB filaments are essential for rod shape when cells adopt nonpolar PG elongation [13, 25, 54].

In stark contrast, in the BN04 cells that expressed both MreB_Ec_-msfCFP^SW^ and MreB_Mx_-PAmCherry^SW^, both MreB homologs remained in polar aggregates in the presence of A22 and cells continued polar growth while maintaining rod shape (**Fig. 3b**). Similarly, when PBP2 was artificially targeted to the poles in the BN10 cells, cells displayed similar resistance against A22, with both MreB_Ec_ and PG elongation focused on cell poles (**Fig. 4a, b**). Taken together, once cells switch their growth sites to cell poles, the Rod enzymes carry out PG elongation independent of MreB filaments.

### Focused PG elongation depends on pre-established cell poles to preserve rod shape

To test whether the MreB protein is still required for focused PG elongation, we acquired the strain ATM1504 that carries an *mreB* deletion but expresses SdiA to suppress *ΔmreB* lethality [55]. The ATM1504 cells grew as spheres and incorporated BADA near homogeneously on their surfaces (**Fig. 4c**). We constructed the BN11 strain to express the PopZ-H3H4-PBP2 fusion *in trans* in the ATM1504 strain with an arabinose-inducible promoter. After 12 h of induction with 0.4% arabinose followed by 15 min of BADA staining, all cells remained spherical.

Two possibilities could account for these observations. First, Rod enzymes may be nonfunctional in the absence of MreB, such that the remaining aPBPs are insufficient to generate or maintain rod shape. Second, PopZ–H3H4–PBP2 does not form stable aggregates without pre-established poles. To test these possibilities, we induced PopZ– H3H4–PBP2 expression in BN11 cells with 0.4% arabinose for 12 h, followed by treatment with moenomycin (4 µg/ ml). After 1 h of inhibition, newly synthesized PG was labeled with BADA for 15 min. Despite inhibition of all aPBPs by moenomycin, induced BN11 cells continued to incorporate BADA (**Fig. 4c**), indicating that Rod enzymes remain active in the absence of MreB_Ec_ and thereby ruling out the first possibility. Thus, without a mechanism that retains Rod enzymes to specific locations, cells cannot initiate focused PG elongation.

## Discussion

Although spherical shapes are energetically efficient and favored by physical principles, growing evidence suggests that the last common ancestor of modern bacteria was rod-shaped [37], and that the evolution of rod-like morphology likely coincided with the emergence of PG [56]. The rigidity of PG is a double-edged sword. It provides sufficient mechanical support for cells but also poses a major hurdle for cell growth and division. Because the entire PG layer is a single molecule, cell growth and division rely on the modification on the existing PG scaffold. To preserve cell width homeostasis while continuously remodeling the cylindrical PG, the elongation machinery in rod-shaped bacteria must fine tune its activity in response to subtle variations in local curvature. Conserved in most rods but absent in most spheres, MreB is an ancient protein that plays critical roles in rod-like morphogenesis [13, 54]. MreB is essential in nearly all rod-shaped bacteria that utilize dispersed, nonpolar elongation because it functions as an effective curvature sensor and dynamic regulator. As MreB filaments move in concert with the Rod polymerases RodA and PBP2, they align along regions of highest membrane curvature, thereby refining the spatial positioning of the elongation machinery [20-22].

In this study, we reprogrammed *E. coli*, the most thoroughly studied rod-shaped bacteria that elongates its PG laterally in a dispersed fashion, to adopt polar elongation—a mode typically restricted to distantly related species. By targeting a single core component of the Rod system—MreB or PBP2—we successfully redirected the entire system to the cell poles, thereby shifting PG elongation to the poles. In this case, as the nonpolar region is no longer the growth zone, the established cylindrical PG structure is sufficient to maintain cell width, where MreB filaments become nonessential. This cross-phylum shift of growth mode highlights the evolutionary plasticity of bacterial morphogenetic systems and suggests that elongation modes, though often conserved within lineages, can be rewired through targeted manipulation of core components.

Due to technical limitations, we cannot exclude the possibility that the reprogrammed, pole-elongating *E. coli* strains still incorporate trace amounts of PG in nonpolar regions—just as we cannot rule out that bacteria growing in nonpolar mode also insert small amounts of PG at the poles. These elongation modes are not necessarily mutually exclusive; rather, individual bacteria may employ a specific blend of both polar and dispersed growth strategies such as recently discovered in *Caulobacteraceae* and *Streptomyces* [3, 4]. However, these two elongation modes may rely on distinct mechanisms to maintain rod shape. Dispersed, nonpolar elongation depends critically on MreB filaments. In *E. coli* and *B. subtilis*, PG-depleted spheroplasts can spontaneously regenerate rod shape through curvature-dependent localization of MreB filaments [21, 57]. Similarly, spherical, PG-free *M. xanthus* spores employ polarity regulators to monitor PG assembly and direct molecular motors that transport MreB filaments—and associated Rod enzymes—away from future poles [35, 36]. By contrast, we propose that in the absence of MreB-mediated modulation, focused PG elongation can sustain rod shape only when cell poles are already established. Without poles, the uniform membrane curvature of spherical cells cannot spatially constrain focused PG synthesis, necessitating alternative polarity determinants such as FtsA, FtsZ, or DivIVA.

An unexpected result is that moenomycin, which inhibits aPBPs, causes MreB_Ec_ filaments to disperse (**Fig. 3a**). This effect is not likely due to the loss of rod shape because MreB_Ec_ still forms filaments in the spherical *ΔrodZ* cells (**Fig. 3d**). Rather than affecting PG directly, moenomycin locks aPBPs in their glycan-charged conformation [58, 59]. Using *M. xanthus*, we discovered that one of the moenomycin-bound aPBPs activates the endopeptidase DacB, which in turn, modifies PG and turns cells into spheres [17]. As endopeptidases are space-makers for the TPases in both PBP2 in the Rod system and aPBPs [1, 60, 61], moenomycin-bound aPBPs could modulate PBP2 in the Rod system through DacB-like endopeptidase in *E. coli*. Such a mechanism also implies that, beyond determining the localization of the entire Rod system [62], PBP2 may also regulate MreB filamentation. In support of this hypothesis, we found that blocking PBP2 directly with mecillinam disperses MreB filament in a similar manner (**Fig. 3a**).

Members of the Rhizobiales and Actinomycetales orders, having lost MreB, maintain rod-like morphology. Strikingly, both orders elongate dominantly from cell poles. Similar to our polar-elongating *E. coli* strains, the established nonpolar regions of these bacteria are seldom modified and thus do not require MreB-like cytoskeleton to maintain cell width homeostasis. Given that MreB is an ancient protein believed to have existed prior to the emergence of *Rhizobiales* and *Actinomycetales* [37], it is plausible that polar growth may have evolved from dispersed elongation as a consequence of MreB loss, which in Rhizobiales and Actinomycetales allows the repurposing of FtsA and FtsZ in the former, and the acquisition of DivIVA in the latter to recruit PG elongation machineries to cell poles [31, 32]. Notably, *Streptomyces venezuelae* can combine MreB-independent polar growth and MreB-dependent dispersed growth [4], which may represent an evolutionary intermediate as reliance on MreB diminished.

## Materials and Methods

**Table S1.**
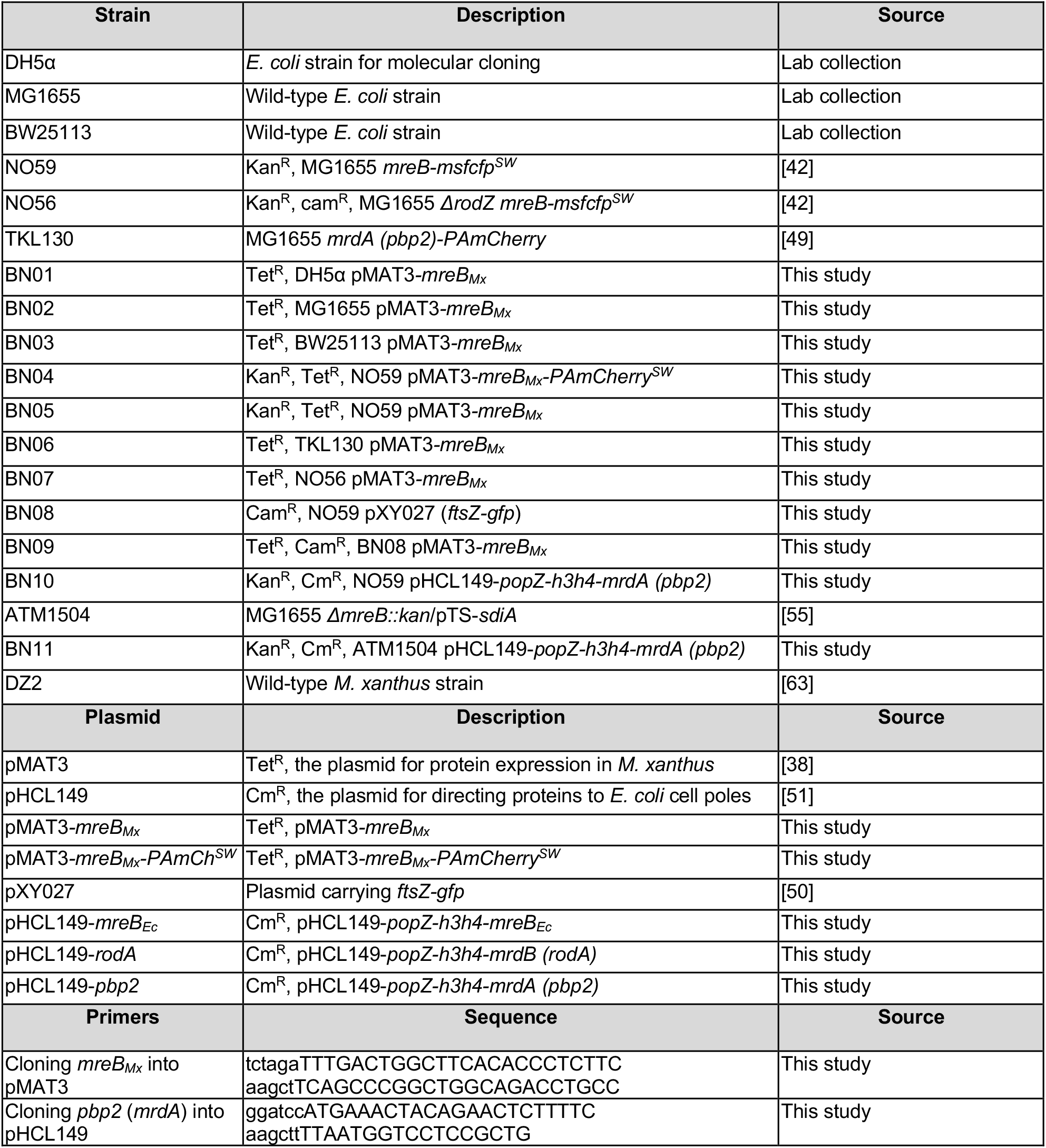
Bacterial strains, plasmids, and primers used in this study.

### Strains and growth conditions

Bacterial strains, plasmids, and primers used in this study are listed in **Table S1**. *E. coli* strains were grown overnight in liquid LB medium with the appropriate antibiotic at 37 °C.

For fluorescent D-amino acid (**FDAA**, including BADA, HADA, and TADA) staining experiments, cells were incubated for 20 min at 37 °C in LB media supplemented with 60 µM FDAA. Cells were collected (9,000 g, 1 min) washed three times with liquid LB and resuspended in ⅕ of the original volume. For sequential staining, cells were incubated with 60 µM of the first FDAA for 15 min, washed three times with liquid LB, incubated with 60 µM the second FDAA for 15 min, and washed three times with liquid LB before imaging. To determine the roles of aPBPs, the Rod system, and MreB in PG elongation, moenomycin (10 µg/ml), mecillinam (100 µg/ml), and A22 (10 µg/ml) was added, respectively, to the culture in liquid LB media 1h before adding FDAA.

### Cryo-EM

*E. coli* cultures were grown in liquid LB medium to OD_600_ ∼1. Cells were collected by centrifugation (3,000 g, 2 min) and resuspended in LB to a final OD_600_ of 12. 3 µl of the cell suspension was applied to a 1.2/1.3 200 mesh copper grid (Quantifoil) that were glow-discharged for 30 s at 15 mA. Grids were plunge-frozen in liquid ethane with an FEI Vitrobot Mark IV (Thermo Fisher Scientific) at 4 °C and 100% humidity, with a waiting time of 30 s, two-side blotting time of 2.5 - 3 s, and blotting force of 0. All subsequent grid handling and transfers were performed in liquid nitrogen. Images were acquired using a Thermo Fisher Scientific Titan Krios G4 transmission electron microscope.

### Live cell imaging

Overnight *E. coli* cultures were diluted to OD_600_ 0.1 in fresh liquid LB medium and grown at 37 °C to OD_600_ 0.4 - 0.6 before imaging. For all imaging experiments, we spotted 5 μl of culture on agar (1.5%) pads. For the treatments with antibiotics, antibiotics were added to both the cell suspension and agar pads. The length and width of cells were determined from differential interference contrast (**DIC**) images using a MATLAB (MathWorks) script [17, 29, 35, 64]. The fluorescence of CFP, BADA, and TADA was excited by different lasers: CFP and HADA at 405 nm, GFP and BADA at 488 nm, and mCherry and TADA at 561 nm. For MreB_Mx_-PAmCherry^SW^ and PBP2-PAmCherry, PAmCherry was activated using a 405-nm laser (0.3 kW/cm^2^, 2 s), excited and imaged using a 561-nm laser (0.2 kW/cm^2^). DIC and fluorescence images of cells were captured using an Andor iXon Ultra 897 EMCCD camera (effective pixel size 160 nm) on an inverted Nikon Eclipse-Ti™ microscope with a 100× 1.49 NA TIRF objective under HILO illumination [29, 35, 45, 46].

### Immunoblotting

The expression of MreB_Mx_ was determined by immunoblotting following SDS-PAGE using a polyclonal anti-MreB_Mx_ serum [40] and a goat anti-Rabbit IgG (H+L) secondary antibody, HRP (Thermo Fisher Scientific, catalog # 31460). The blots were developed with Pierce™ ECL Western Blotting Substrate (Thermo Fisher Scientific REF 32109) and a MINI-MED 90 processor (AFP Manufacturing).

## Acknowledgments

We thank Drs. Michael VanNieuwenhze and Yen-Pang Hsu for providing TADA and HADA, Drs. Randy Morgenstein, KC Huang, and Anthony Maurelli for sharing the strains NO59, NO56, TKL130, and ATM1504 and Dr. Gaya Yadav for the assistance on cryoEM imaging. Part of this work was supported by the National Institutes of Health grants GM129000 to B. N.. We received financial support from Dr. David R. Zusman, who played no role in the design, execution, or presentation of this work.

---

**Fig. S1.**
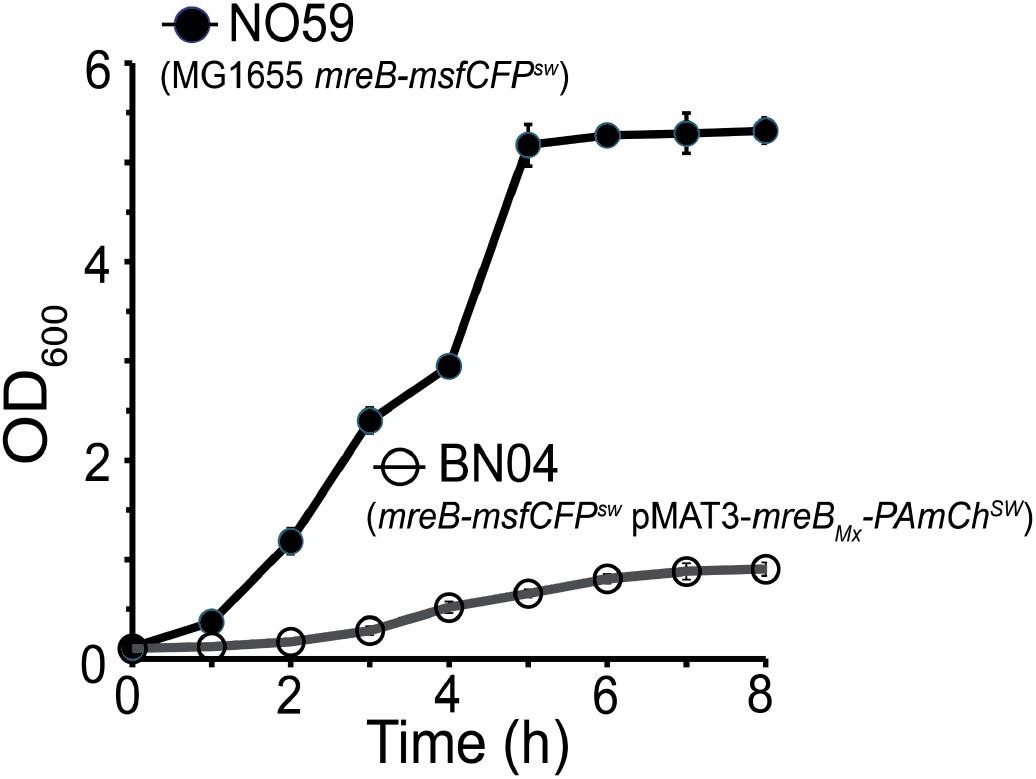
BN04 cells grow slower than its parental strain NO59. Averages and standard deviations were calculated from three technical repeats.

## Notes

### Competing Interest Statement

The authors have declared no competing interest.

